# Diversity and Life-cycle Analysis of Pacific Ocean Zooplankton by Videomicroscopy and DNA Barcoding. I. Cnidaria

**DOI:** 10.1101/669663

**Authors:** Peter J. Bryant, Timothy E. Arehart

**Affiliations:** Department of Developmental and Cell Biology, University of California, Irvine, Irvine, CA 92697; Crystal Cove Conservancy, #5 Crystal Cove, Newport Coast, CA 92657

## Abstract

Most, but not all cnidarian species in the classes Hydrozoa, Scyphozoa and Anthozoa have a life cycle in which a colonial, asexually reproducing hydroid phase alternates with a free-swimming, sexually reproducing medusa phase that, in the hydrozoans, is usually microscopic. Hydrozoan medusae were collected by zooplankton tows in Newport Bay and the Pacific Ocean near Newport Beach, California, and hydroid colonies were collected from solid substrates in the same areas. Specimens were documented by videomicroscopy, preserved in ethanol, and sent to the Canadian Centre for DNA Barcoding at the University of Guelph, Ontario, Canada for DNA barcoding.

Among the order Anthomedusae (athecate hydroids), DNA barcoding allowed for the discrimination between the medusae of eight putative species of *Bougainvillia*, and the hydroid stages were documented for two of these. The medusae of three putative species of *Amphinema* were identified, and the hydroid stages were identified for two of them. DNA barcodes were obtained from medusae of one species of *Cladonema*, one adult of the By-the wind Sailor, *Velella Velella*, five putative species of *Corymorpha* with the matching hydroid phase for one; and *Coryne eximia, Turritopsis dohrnii* and *Turritopsis nutricula* with the corresponding hydroid phases. The actinula larvae and hydroid for the pink-hearted hydroid *Ectopleura crocea* were identified and linked by DNA barcoding.

Among the order Leptomedusae (thecate hydroids) medusae were identified for *Clytia elsaeoswaldae, Clytia gracilis* and *Clytia sp. 701 AC* and matched with the hydroid phases for the latter two species. Medusae were matched with the hydroid phases for two species of *Obelia* (including *O. dichotoma*) and *Eucheilota bakeri. Obelia geniculata* was collected as a single hydroid. DNA barcodes were obtained for hydroids of *Orthopyxis everta* and three other species of *Orthopyxis*.

The medusa of one member of the family Solmarisidae, representing the order Narcomedusae, and one member (*Liriope tetraphylla*) of the order Trachymedusae were recognized as medusae.

In the Scyphozoa, DNA barcoding confirmed the planktonic larval stage (ephyra) of the Moon Jelly, *Aurelia aurita*, the adult medusa of which is occasionally common in and around Newport Bay. In the Anthozoa, antipathula larvae were identified from the Onion Anemone, *Paranthus rapiformis* and a cerinula larva was identified from the Tube-dwelling Anemone, *Isarachnanthus nocturnus*. We have yet to find the adults of these species locally.

## Introduction

Many cnidarian species alternate between two related body forms - a sexually reproducing, free-swimming medusa phase and an asexually reproducing hydroid stage, which often exists as a sessile colony. Both body forms show a basic radial symmetry, with a mouth surrounded by tentacles leading into the body cavity where digestion occurs. In some species, either the medusa or the hydroid phase is missing. Two of the cnidarian classes – the Scyphozoa (Sea Jellies) and Anthozoa (Sea Anemones) are well known, but the class Hydrozoa is less well known, in part because many of the species are microscopic at least in the medusa phase. In some cases the medusa and hydroid phases have been described as separate species, and matching the two phases would have required rearing of the organism from one phase to another, which has not always been possible. Here we show that DNA barcoding makes it possible to easily link life cycle phases without the need for laboratory rearing. The results also provide preliminary data on the extraordinary level of biodiversity within the Hydrozoa.

## Methods

Zooplankton collections were made from the following sites under Scientific Collecting Permit SC-12162 from the California Department of Fish and Wildlife:

| Locality | Latitude | Longitude |
| --- | --- | --- |
| Balboa Island Coral Street | 33D 36'N | 117D 53'W |
| Balboa Pavilion Dock | 33D 36'N | 117D 54'W |
| Bayside Marina | 33D 37'N | 117D 54'W |
| Back Bay Science Center Dock | 33D 37'N | 117D 53'W |
| Cabrillo Marine Aquarium | 33D 43'N | 118D 17'W |
| Crew dock, BBSC | 33D 37'N | 117D 53'W |
| Delhi Channel | 33D 38'N | 117D 53'W |
| Lido Island (Via lido and Genoa) | 33D 36'N | 117D 55'W |
| Mussel Rocks, Crystal Cove State Park | 33D 34'N | 117D 50'W |
| Newport Harbor | 33D 36'N | 117D 53' W |
| Newport Harbor Entrance | 33D 35'N | 117D 53'W |
| Newport Pier | 33D 36'N | 117D 55'W |
| Ocean off Crystal Cove State Park | 33D 33'N | 117D 51'W |
| Ocean off Dana Point | 33D 27'N | 117D 42'W |
| Ocean off Newport Beach | 33D 34'N | 117°53' W |
| Off Newport Aquatic Center | 33D 37'N | 117D 53'W |
| Under Pacific Coast Highway Bridge | 33D 37'N | 117D 54'W |
| Reef Point, Crystal Cove State Park | 33D 34'N | 117D 50'W |
| San Clemente Beach | 33D 25'N | 117D 37'W |

Collections were made with a 150 µm mesh net attached to a rope, with a 50 ml collection tube at the base. They were made from a public dock using repeated horizontal sweeps near the surface and diagonal sweeps down to about 15 ft. depth. About 5-10 sweeps of a total of about 100 feet usually yielded sufficient specimens, but no attempt was made to monitor collections quantitatively. Collections were made and analyzed with the assistance of Undergraduate students Taylor Sais, Alicia Navarro, Debbie Chung and Lesly Ortiz.

Ocean collections were made with a 150 micron mesh net attached to a 100 ft. rope. The net (aperture 30 cm) was towed behind the vessel, just below the surface, for a period of 7 minutes at the slowest possible speed. Deployment and retrieval extended the total tow period to 10 minutes.

Collections were brought to the laboratory at the University of California, Irvine and examined under a dissecting microscope with lateral light and a dark background. Each specimen of interest was removed using a Pasteur Pipette, transferred to a depression slide, and recorded by video microscopy using a Zeiss microscope with a dark-field condenser, fitted with a phototube attached to a Nikon 5100 single-lens reflex camera. The most informative frames were taken from the videos and used in the figures for this paper. Each specimen was preserved in 90% ethanol in a well of a 96-well microplate. Filled plates were sent to the Canadian Centre for DNA Barcoding at the University of Guelph for sequencing of the standard 648-bp “DNA barcode” (Hubert and Hanner, 2015) in the COI mitochondrial gene, using the following primers: (LCO1490 GGTCAACAAATCATAAAGATATTGG; HCO2198 TAAACTTCAGGGTGACCAAAAAATCA). This usually produced a DNA barcode of 658 nucleotides, and only those containing >/= 600 nucleotides were included in the sequence analysis. Species that could not be identified morphologically were assigned operational taxonomic names of the form nPJB, and groups of specimens with identical or almost identical DNA barcodes were assigned BIN numbers. The DNA sequences are in the public domain at the Canadian Centre for DNA Barcoding.

Hydroid phase specimens were obtained by removing bunches of seaweed from docks in Newport Bay, bringing them to the laboratory and examining them under the dissecting microscope. This provided hydroid stages for many but not all of the medusae investigated, presumably because hydroids of some species live on the seabed or in other locations that are difficult to access for collections. Hydroid specimens were recorded and processed in the same manner as medusae.

## Results

From 843 Specimens, 497 sequences were obtained falling into 97 BINs, containing 39 recognized species. The specimen growth curve shows that this collection is approaching saturation, but has not yet reached it:

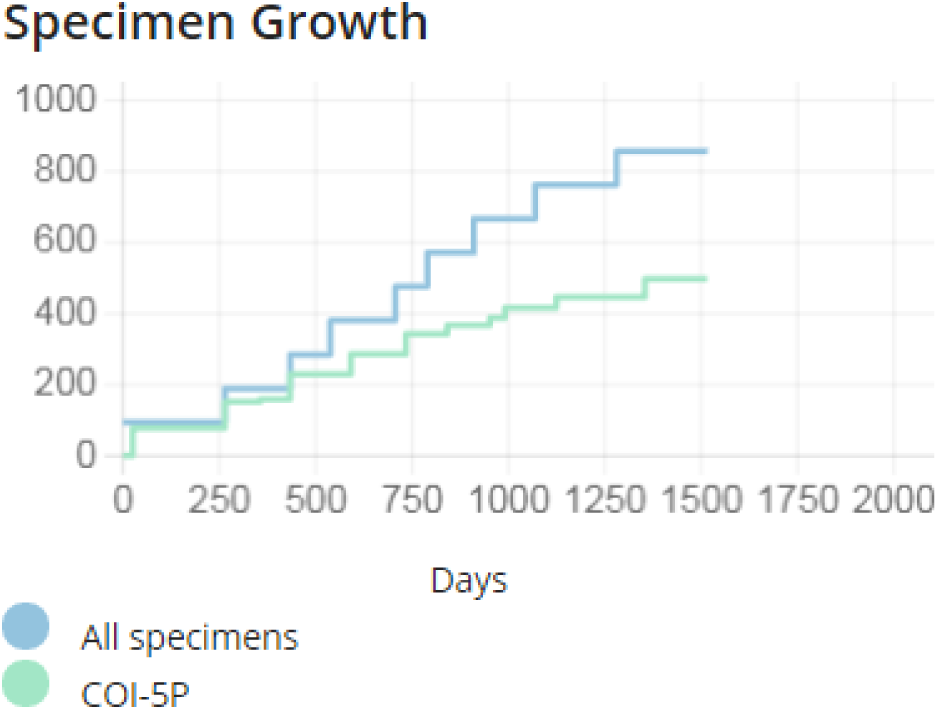

Cnidarians are divided into four classes (Hydrozoa, Scyphozoa, Anthozoa, Cubozoa), of which the first three are represented in the present collection. The fourth Class, <u>Cubozoa</u> (box jellies) is recognized and has been documented off Southern California by a single species collection in San Diego County (Straehler-Pohl et al., 2017).

### Class Hydrozoa

These usually have the typical cnidarian life cycle, alternating between sexually reproducing medusae and asexually reproducing hydroids. The hydroids are typically in colonies, which can be either male or female.

* Representatives found in the current collection.

Subclass Leptolinae

* Order Anthomedusae (Anthoathecata; with athecate hydroids)
* Order Leptomedusae (Leptothecata; with thecate hydroids)
* Order Siphonophorae

Subclass Trachylinae

Order Actinulidae

Order Limnomedusae

* Order Narcomedusae
* Order Trachymedusae

#### Order Anthomedusae: Athecate Hydroids

The hydroid stalk is naked (athecate; not surrounded by a sheath).

##### Family Bougainvilliidae

Medusae of local species in this family are all microscopic and can be distinguished by the number of tentacle bases and the number of tentacles attached to each base (Vannucci and Rees, 1961). The digestive system hangs from the center of the umbrella, leading to the manubrium with the mouth at its tip. Some species also have four small tentacles surrounding the mouth. The gonads, producing sperm in the males and eggs in the females, are attached to the sides of the manubrium.

The hydroid stock is athecate, the hydranths having a conical proboscis and a single whorl of filiform tentacles; the medusa buds are borne below the hydranths, on their pedicels or on the stems. The hydroids in this family are very difficult to identify because they show much less obvious morphological diversity than the corresponding medusa phases.

###### Genus *Bougainvillia*

Medusa with four radial canals and four single or bundled marginal tentacles; gonads on the side of the manubrium (Vannucci and Rees, 1961; Schuchert, 2007) (Figs 1 and 2).

**Fig 1.**
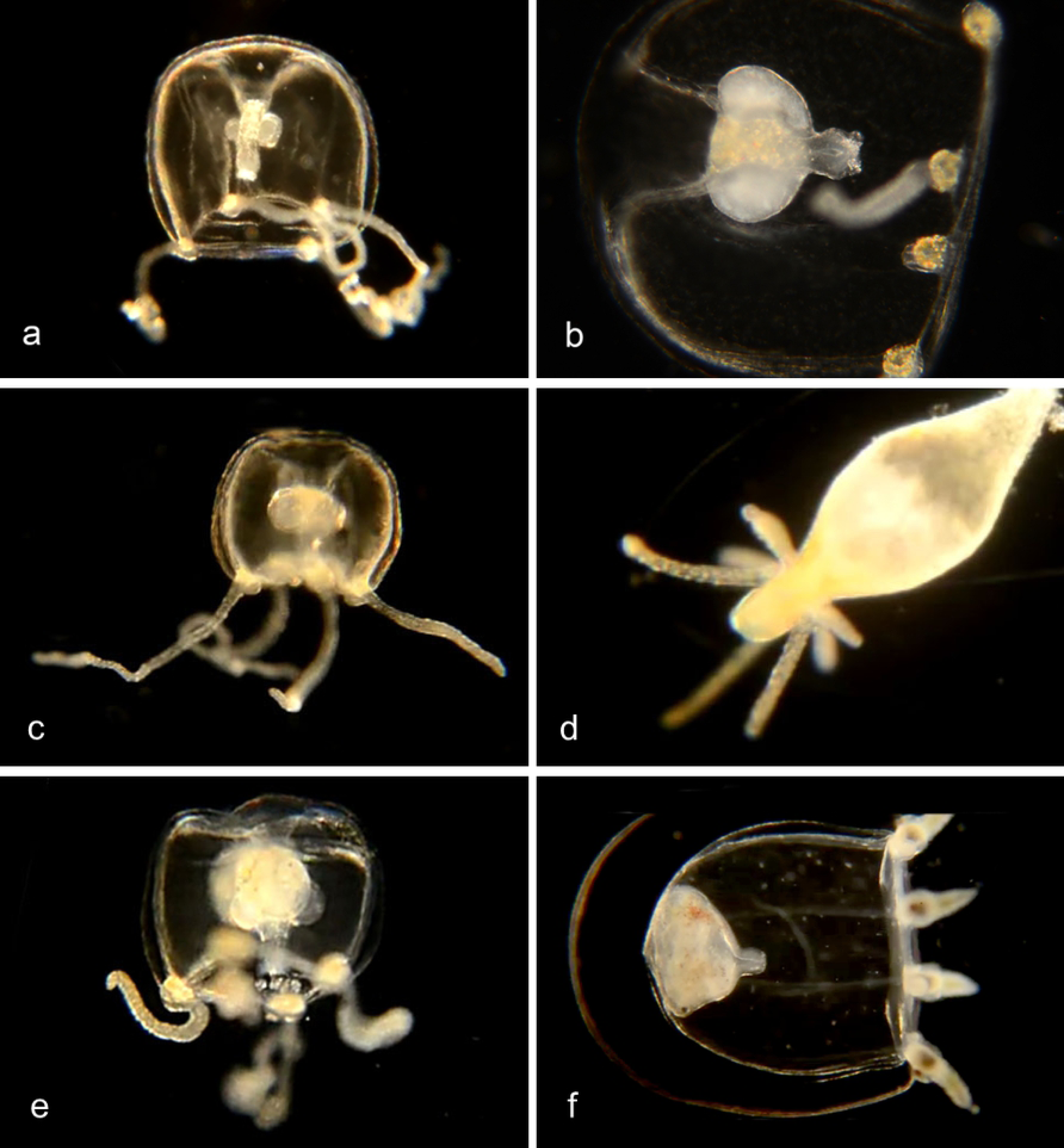
Species with four single tentacles. a,b. *Bougainvillia sp*. 1PJB (Bin ACR4341): Medusa with four radial canals and four single tentacles; oral tentacles present. Material collected: 16 medusae (11 from Newport Pier, 2 from Balboa at Coral, 3 from Ocean off Dana Point). a, Medusa BIOUG01227-E08. b, Medusa BIOUG01213-C03. c,d,e. *Bougainvillia sp*. 2PJB (BIN ACR4343): Medusa with four radial canals and four single tentacles, and four oral tentacles. Material collected: 27 medusae (10 off Newport Pier, 12 off Balboa at Coral, one in Delhi Channel, 2 in Ocean off Crystal Cove, one in Ocean off Newport, one in Harbor entrance), one hydroid off Newport Pier. c, Medusa BIOUG01227-D05. d, Hydroid BIOUG01227-A07. e, Manubrium with medusa buds and oral tentacles BIOUG19284 H06; f, *Bougainvillia sp*. 3PJB (BIN ACW4756): Four radial canals, four unbranched tentacles, no oral tentacles. Material collected: One medusa BIOUG01227-G02, from Newport Pier.

**Fig 2.**
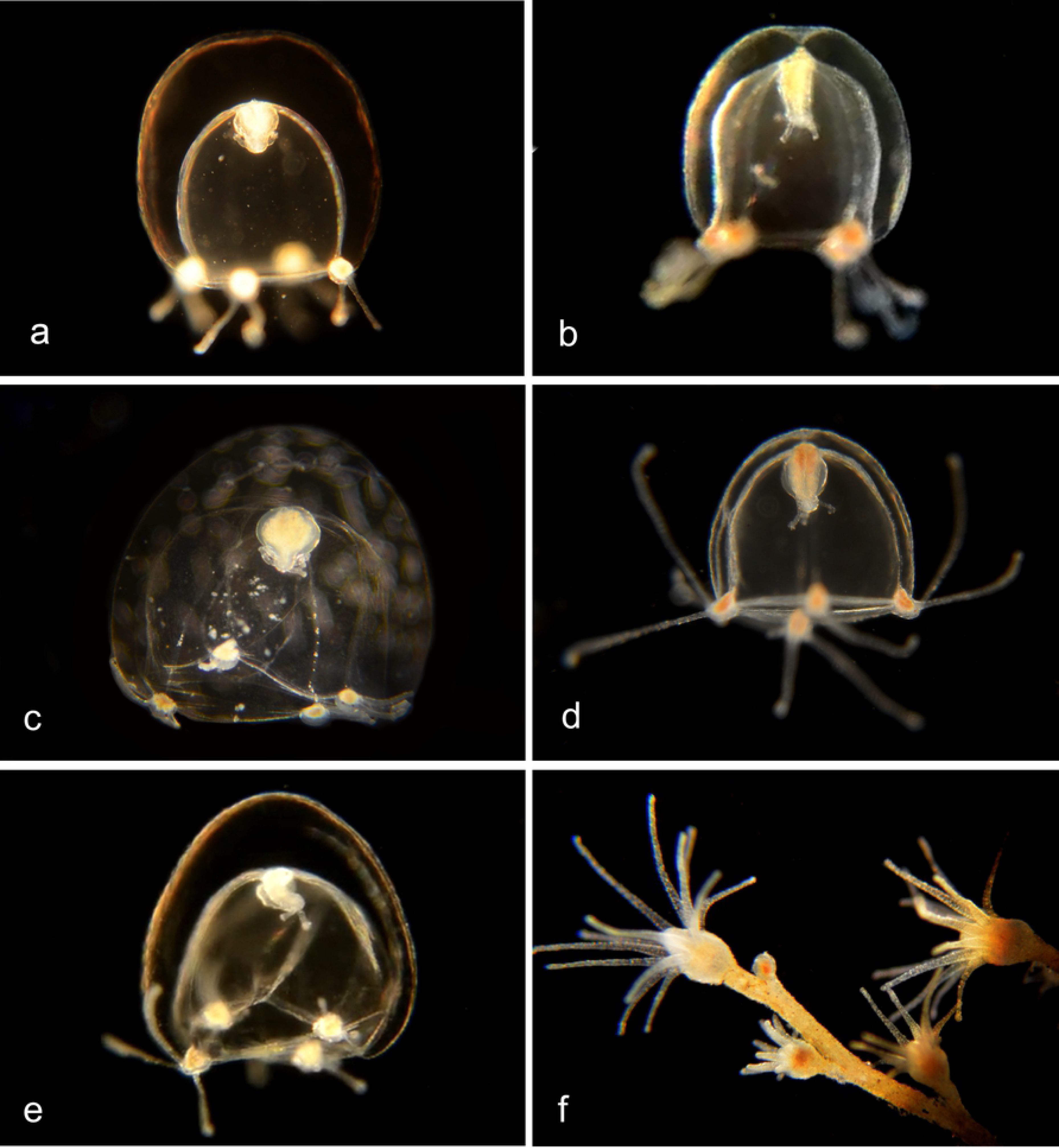
Species with four pairs or triplets of tentacles. a. *Bougainvillia muscus* (BIN ACO8375). Medusa with four radial canals, four pairs of tentacles and four oral tentacles. Material collected: Two medusae, from Lido Island. These show a close match (>98%) by DNA barcode with specimens of *Bougainvillia muscus* collected in Skagerrak (Sweden), Roscoff (France) and New Brunswick (Canada). BIOUG01213-H09. b. *Bougainvillia sp*. 4PJB (BIN ACR4338). Medusa with four radial canals, four pairs of tentacles and four oral tentacles. Material collected: 7 medusae, all from off Newport Pier. BIOUG01213-G06. c. *Bougainvillia sp*. 5PJB (BIN ADC0389). Medusa with four radial canals, four pairs of tentacles and four oral tentacles. Material collected: One medusa (only 607 nucl.), from Balboa at Coral. BIOUG19285-A03. d. *Bougainvillia sp*. 6PJB (BIN ACR 4342). Medusa with four radial canals, four triplets of tentacles and four oral tentacles. Material collected: Three medusae (1 from Newport Pier, two from Lido Island) BIOUG01227-C08. e,f. *Bougainvillia sp*. 7PJB (BIN ACR 4339): Medusa with four radial canals, four triplets of tentacles, two oral tentacles. Material collected: One medusa (Crew Dock), one hydroid colony (Newport Aquatic Center). a, Medusa BIOUG01213-D09; b, Hydroid BIOUG19284-F02.

###### Genus *Amphinema*

Medusa with four radial canals and two single tentacles, no oral tentacles; Hydroids with a conical proboscis and a single whorl of simple tentacles (Schuchert, 2007) (Fig 3).

**Fig 3.**
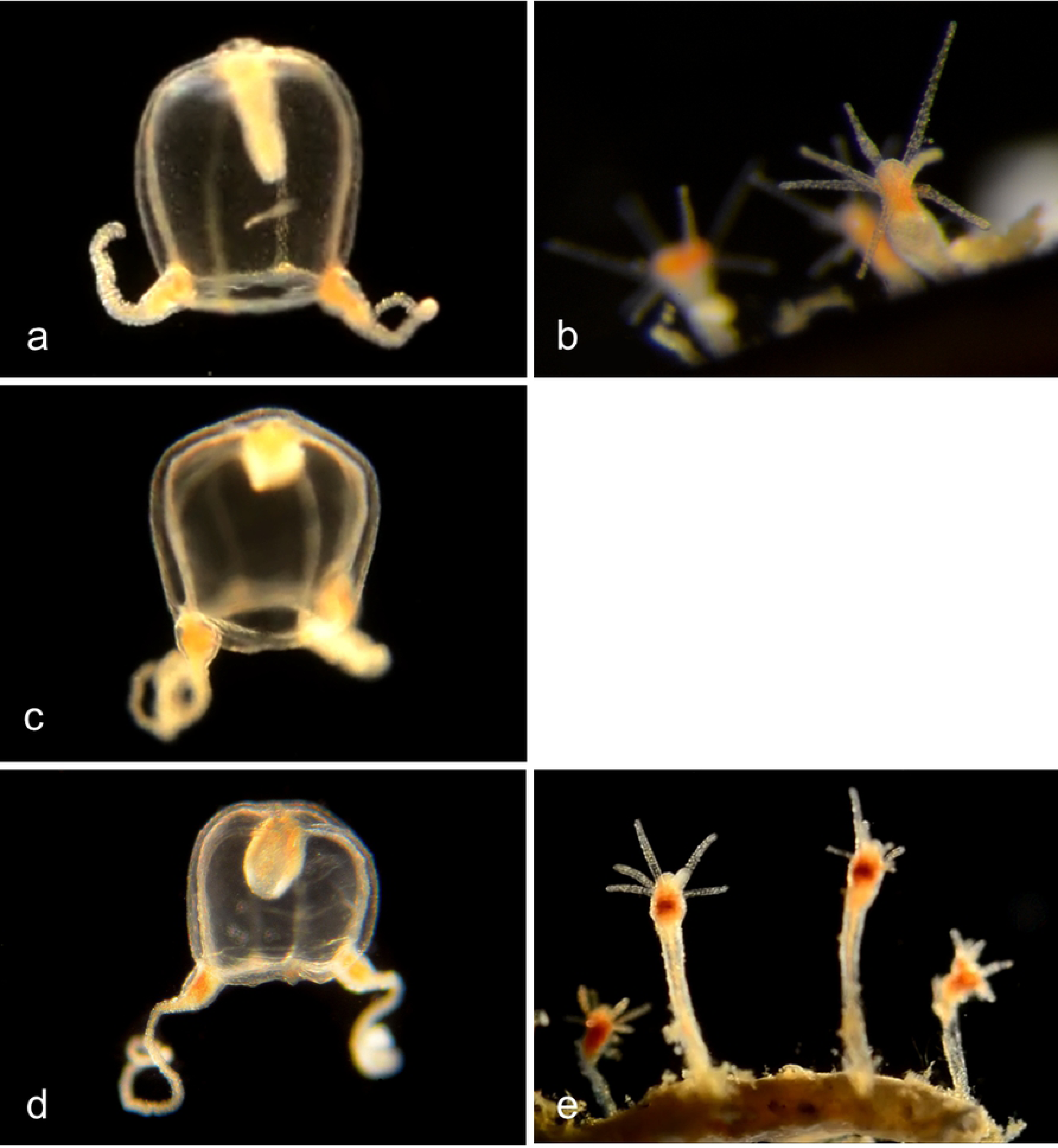
*Amphinema* species. a,b. *Amphinema sp*. 1PJB (BIN ACR4255): Material collected: four medusae (3 from Newport Pier, one from Harbor entrance), three hydroid colonies, all from Balboa Pavilion dock. a, Medusa BIOUG19284-G12; b, Hydroid BIOUG19285-D12. c. *Amphinema dinema* (BIN ACW2490): Material collected: One medusa, from Newport Pier. 100% match of DNA barcode COI with a specimen from Roscoff, France (Schuchert, 2007). Medusa BIOUG01227-H02. d,e. *Amphinema sp*. 2PJB (BIN ACR4184): Material collected: Four medusae (Two from Balboa Pavilion Dock, one from Bayside Marina, one from Newport Pier), three hydroid colonies (two from Newport Pier, one from Balboa Pavilion Dock) d, Medusa BIOUG01227-H10; e, Hydroid BIOUG19285-D07.

##### Family Cladonematidae

The family Cladonematidae, represented by the genus Cladonema, is sometimes collected in plankton samples, but its habitat is reported to be on the leaves of eelgrass, where the inner branches of all nine tentacles are used as suckers and the outer branches carry stinging cells. At the base of each tentacle is a red ocellus (eye-like structure) (Hyman, 1947; Schuchert, 2006). Material collected (Fig. 4; BIN ACR4340): 7 medusae (3 at Lido Island; two at Balboa at Coral, one at Back Bay Science Center Dock, one at Crew Dock).

**Fig 4.**
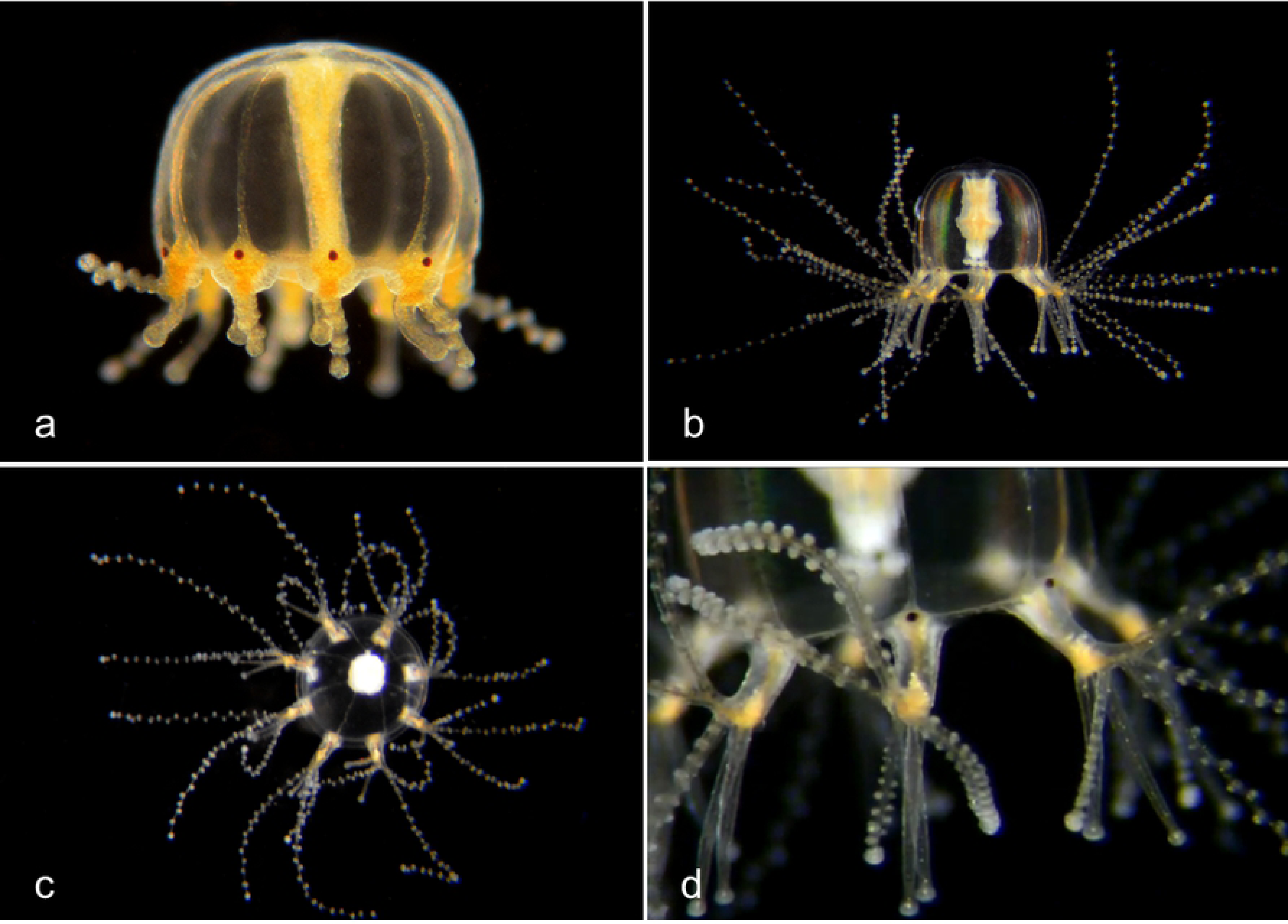
*Cladonema californica*. a, Medusa BIOUG19284-C05. b, c, d, This specimen was found in tanks at Orange Coast College, Orange County, CA on 7/19/2018. It probably arrived in a shipment of sea urchins collected in Long Beach, CA. Hydroid stage not yet found locally.

##### Family Corymorphidae

In the genus *Corymorpha* (Schuchert, 2010) the medusa is distinguished by having one of the four tentacles much larger than the others and containing most or all of the stinging cells. Four radial canals, No oral tentacles. The hydroid has one ring of tentacles around the mouth and another ring more basally (Fig 5).

**Fig 5.**
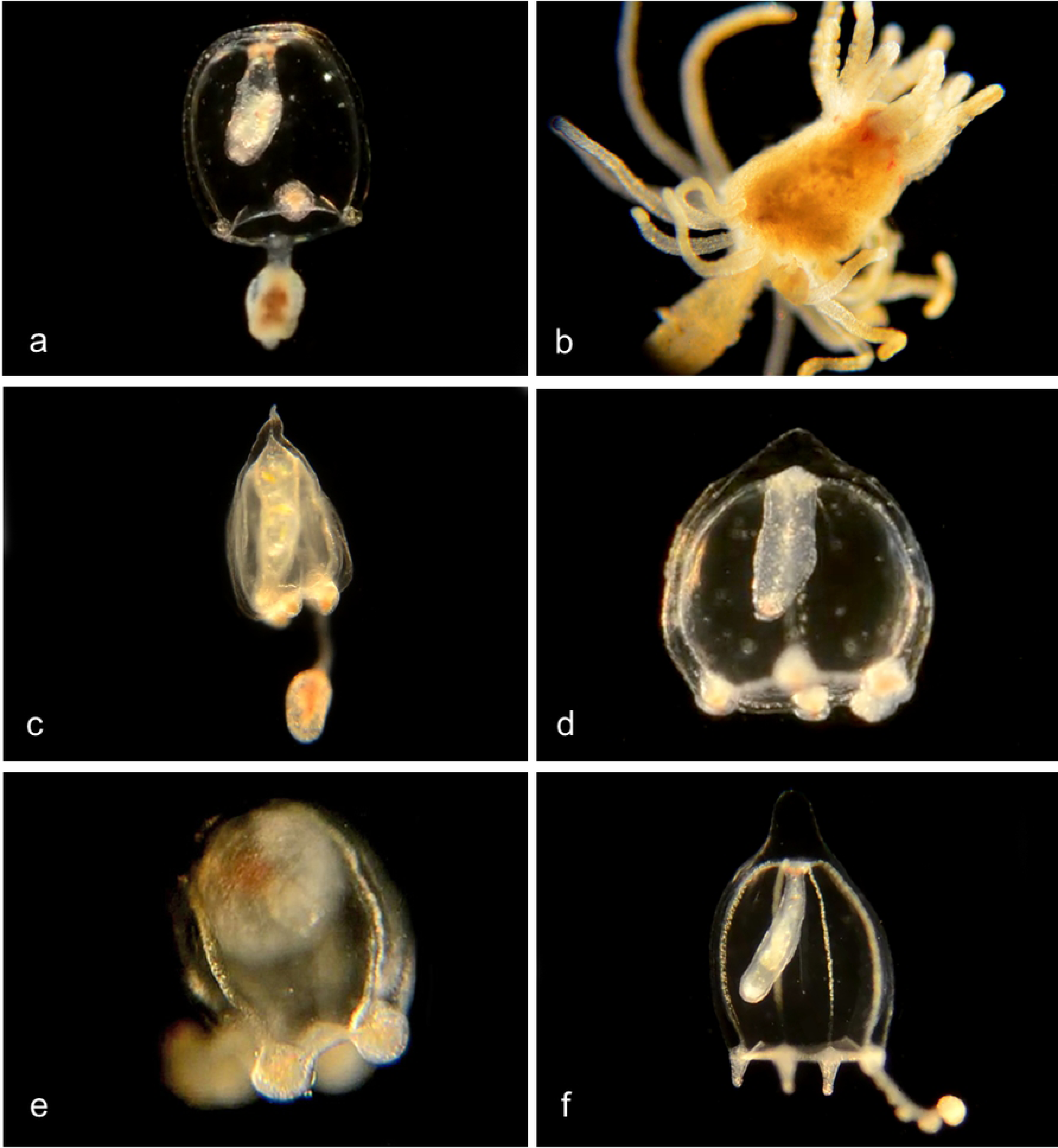
*Corymorpha* species. a, b. *Corymorpha sp. 1PJB* (BIN ACR4296). Material collected: 13 medusae (3 at Crew Dock, 3 at Balboa at Coral, 3 at Lido, and one each at NAC, Delhi channel, Newport Pier and Bayside Marina); two hydroids (one at Balboa at Coral, one at Crew Dock). a, Medusa BIOUG19284-H03; b, Hydroid BIOUG19289-A08. c.*Corymorpha sp. 2PJB* (BIN ADO3522). Material collected: 1 medusa, at Harbor entrance. Medusa CCDB24005-H02. d. *Corymorpha sp. 3PJB* (BIN ACZ1499). Material collected: 3 medusae (1 at Lido at Genoa, 1 at Crew Dock, 1 at Bayside Marina). Medusa BIOUG19284-E04. e. *Corymorpha sp. 4PJB* (BIN ACH5005) close DNA match to both *C. bigelowi* and *C. verrucosa*. Material collected: 1 medusa (at Lido at Genoa). Medusa BIOUG01213-G01. f. *Corymorpha bigelowi* (BIN ACH5002). (Sassaman and Rees, 1978). Material collected: 21 medusae (8 from NAC, 4 from Delhi Channel, 4 from Crew Dock, 2 from Newport Pier, 2 from Lido Island at Genoa, and 1 from Balboa Island at Coral), no hydroids. Medusa BIOUG19284-H07.

##### Family Corynidae

Genus *Coryne*. Medusa has four radial canals and four tentacles with nematocysts concentrated at the tips; long manubrium with no oral tentacles. Stolon is branched, hydroids club-shaped with scattered knob-shaped tentacles carrying clusters of nematocysts at their tips (Fig 6) (Schuchert, 2001).

**Fig 6.**
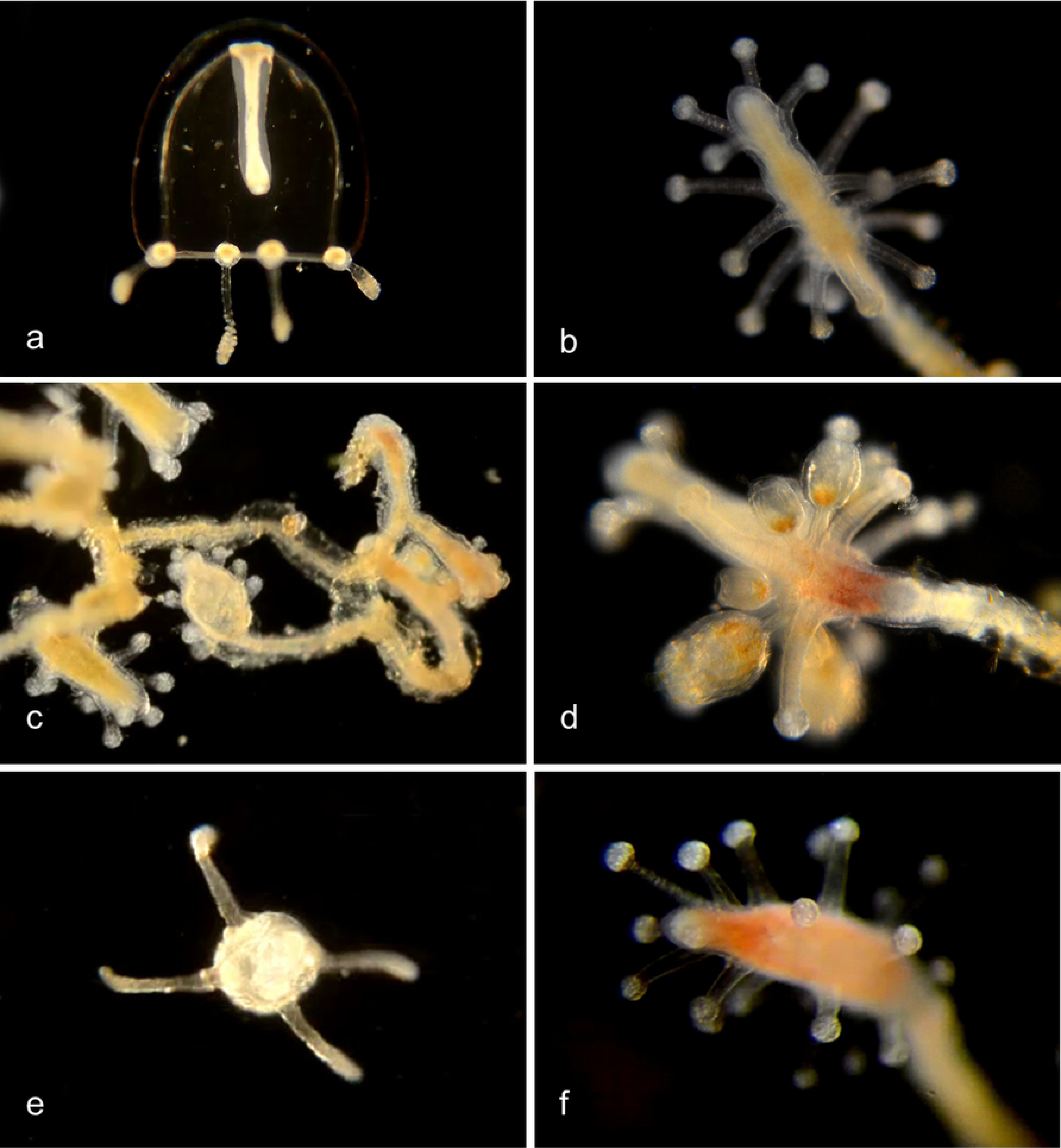
*Coryne* species. a-e. *Coryne eximia* (BIN ACR4295). Material collected: 17 medusae (9 at Newport Pier, 5 at Balboa at Coral, 1 each at Crew Dock, Balboa Pavilion dock, and Reef Point CCSP), 8 hydroids (4 at Newport Pier, 2 at CCSP and 1 each at Mussel Rocks and Reef Point, CCSP) and one actinula larva from Newport Pier. These 26 specimens show closely matched DNA barcodes with one another but with no other sequences in BOLD or Genbank. One other (CCDB 24005 C11 from Balboa at Coral) shows a closer match with one specimen from Norway and one from South Africa, and one more (BIOUG01213 F05 from Newport Pier) shows a closer match with a specimen from Chile. a, Medusa BIOUG01227-A09. b, Club Hydroid BIOUG19289-E06. c, Club hydroid BIOUG19289 F01. d, Club hydroid BIOUG19289-E05. The medusa buds arise in the tentacle axils. e, Actinula larva BIOUG01227-H04. f, *Coryne uchidai* (BIN ACR4087) Club hydroid BIOUG01213-G02: Material collected: 1 hydroid at Lido at Genoa. DNA barcode shows 98.77% match with a specimen from Hokkaido, Oshoro, Japan.

##### Family Oceaniidae

Genus *Turritopsis* (Fig 7)(Schuchert, 2004). Medusa with 4 radial canals and 8-12 tentacles (80-90 in literature); manubrium with no oral tentacles. Filiform tentacles scattered over much of the hydranth body.

**Fig 7.**
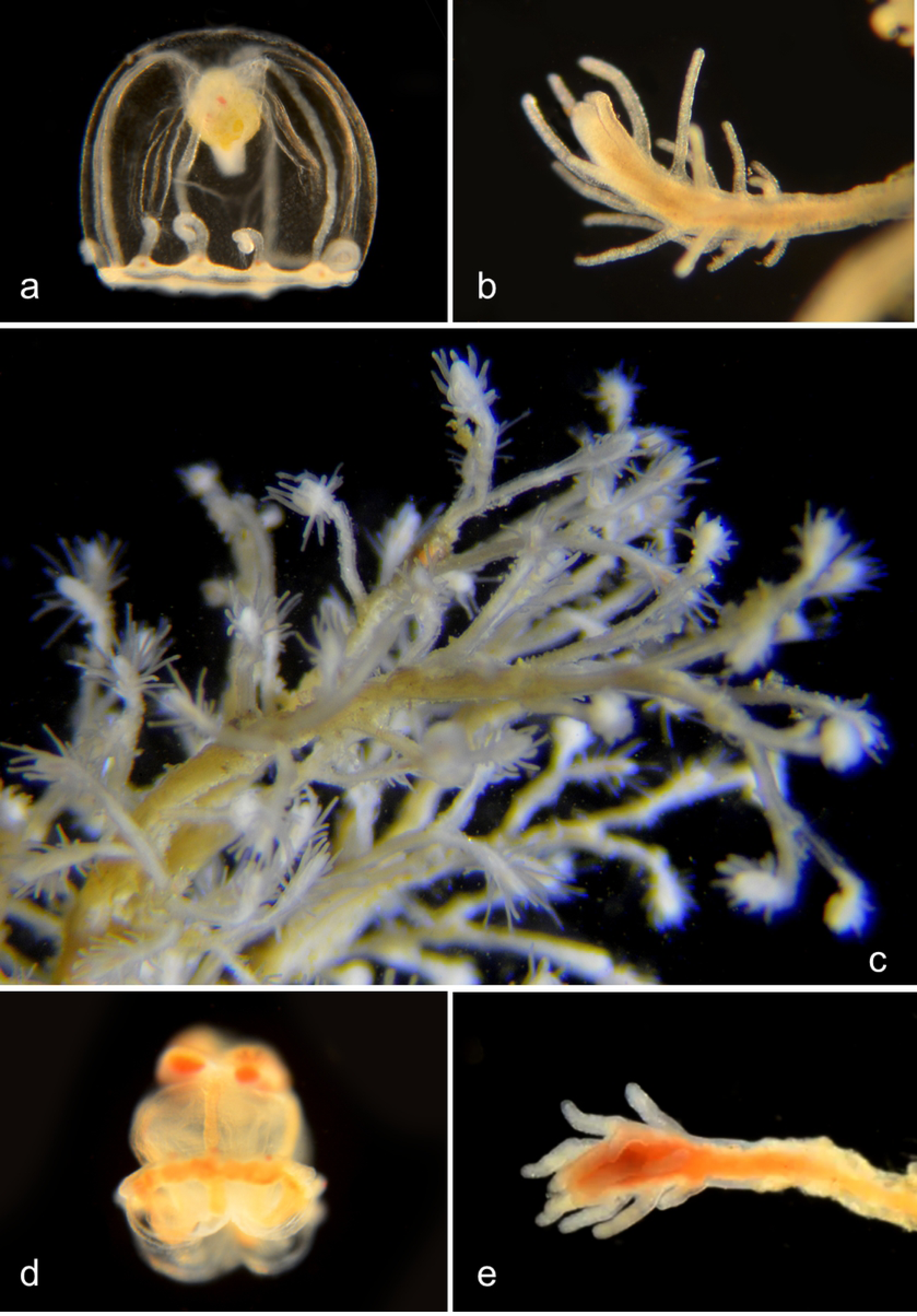
*a-c. Turritopsis dornii* (BIN ACR4293). Material collected: 8 medusae (2 from NAC, 2 from Balboa/Coral, 3 from Lido/Genoa, one from Balboa Pavilion); one hydroid (from Pavilion Dock). a, Medusa BIOUG01227-C01; b, Club hydroid BIOUG19285-E09; c, Club hydroid BIOUG19285-E09. d,e. *Turritopsis nutricula* (BIN ACI0369). Material collected: d, One medusa (inverted?) BIOUG19285-G08 (From Newport Harbor entrance); e, Club hydroid CCDB-25431-F11 (From Newport Pier).

##### Family Porpitidae

By-the wind Sailor, *Velella Velella* (BIN ACR4084) (Fig 8). (Brinckmann-Voss, 1970). Each of these floating jellies is a colony, composed of many individuals that are specialized for various functions: the gonozooids carry out feeding and reproduction, and the dactylozooids protect the colony using stinging cells. The stiff sail catches the wind and propels the colony at a slight angle from directly downwind over the ocean surface. Under some conditions, thousands of them can wash ashore on beaches.

**Fig 8.**
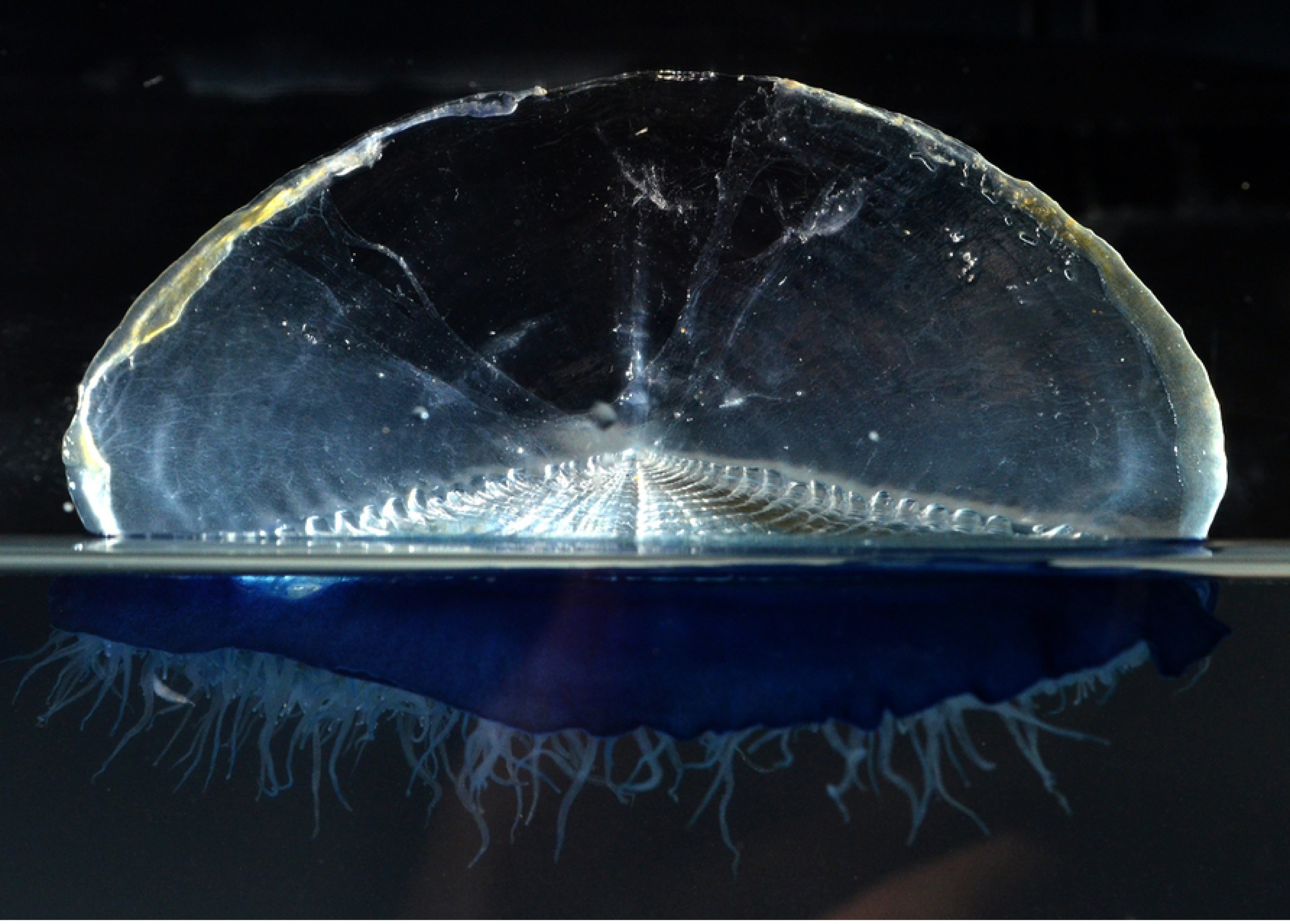
By-the wind Sailor, *Velella Velella*. Material collected: One colony, from San Clemente Beach.

##### Family Tubulariidae

Pink-hearted hydroid, *Ectopleura crocea* (BIN ACH9225) (Fig 9). (Fofonoff et al., 2019, Schuchert, 2010). Exotic species, native to the east coast of North America. It lacks a medusa stage, but grows as colonies of hydroids, each of which has two whorls of tentacles. Medusa buds (“medusoids”) develop between the two whorls of tentacles, but are not released. They produce either eggs or sperm, and internal fertilization occurs in the female medusoids. The fertilized eggs develop into actinula larvae, which are released and develop directly into hydroids. Material collected: 3 actinula larvae (2 from Newport Pier, one from NAC), 5 hydroids (2 from Newport Pier, one from Balboa/Coral, one from Ocean, one from Harbor entrance).

**Fig 9.**
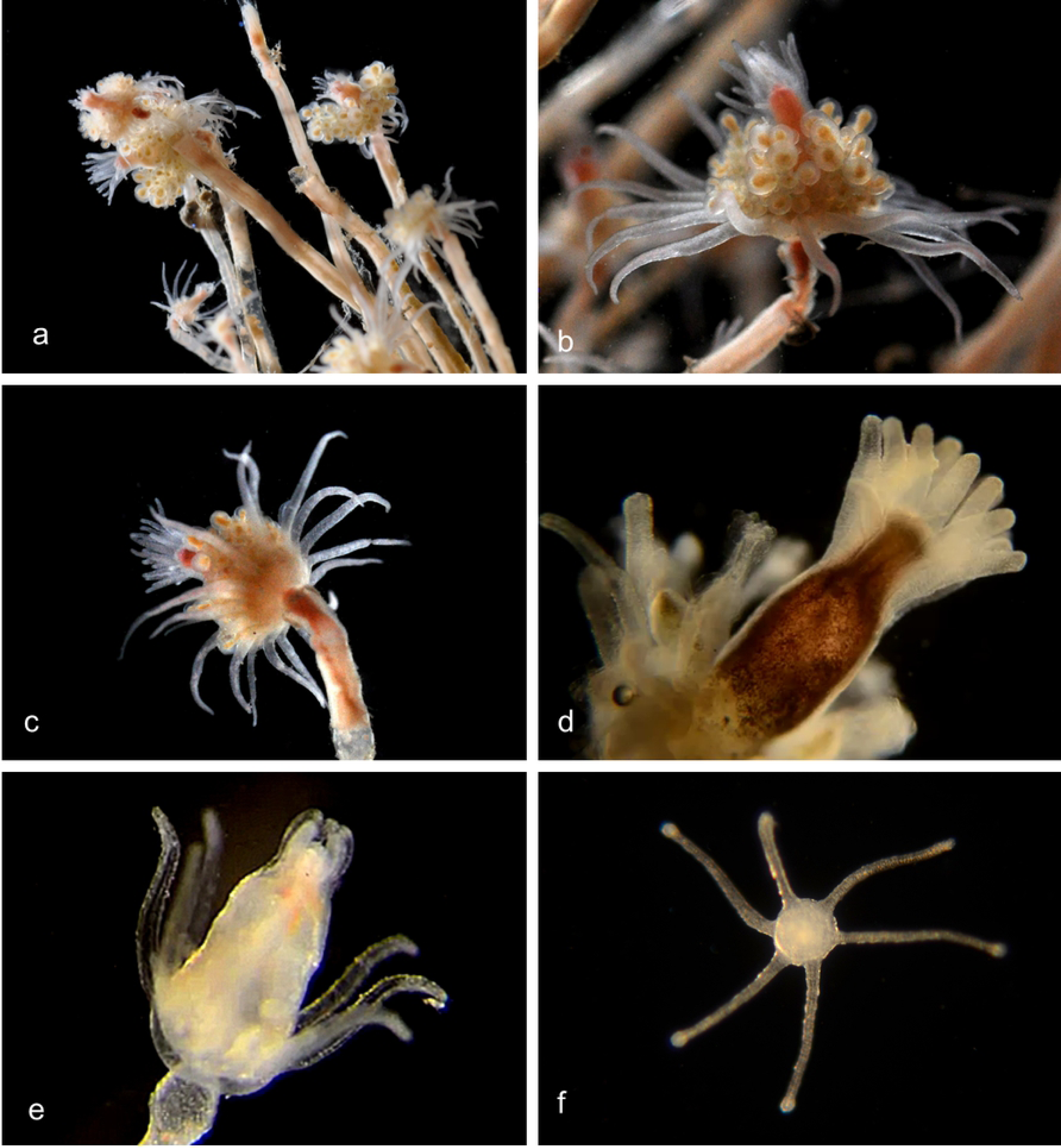
Pink-hearted hydroid, *Ectopleura crocea*. a, Hydroid colony (Not barcoded); b, Hydroid (Not barcoded); c, Hydroid (Not barcoded); d, Hydroid BIOUG01227-C07; e, Hydroid BIOUG01213-G03; f, Actinula larva BIOUG01213-E05.

#### Order Leptomedusae: Thecate hydroids

These have both medusa and hydroid stages, and the medusae are usually larger than in the Anthomedusae. The hydroid stalk is thecate (surrounded by an acellular sheath). The medusae of various species are difficult or impossible to distinguish from one another, so the taxonomy is based on the hydroid phase (the opposite to the situation in Anthomedusae).

##### Family Campanulariidae

Notable genera are *Clytia, Obelia* and *Orthopyxis* (Cornelius, 1982). The hydroid colony reproduces asexually. During the hydroid stage of the life cycle, colonies are attached to substrate surfaces. A mature colony carries feeding hydroids called gastrozooids, defensive hydroids, and reproductive hydroids (gonozooids), which produce medusae by budding. The umbrella-shaped medusa has four or more unbranched tentacles, and the gonads are on the radial canals.

###### Genus *Clytia* (He et al., 2015) (Fig 10)

**Fig 10.**
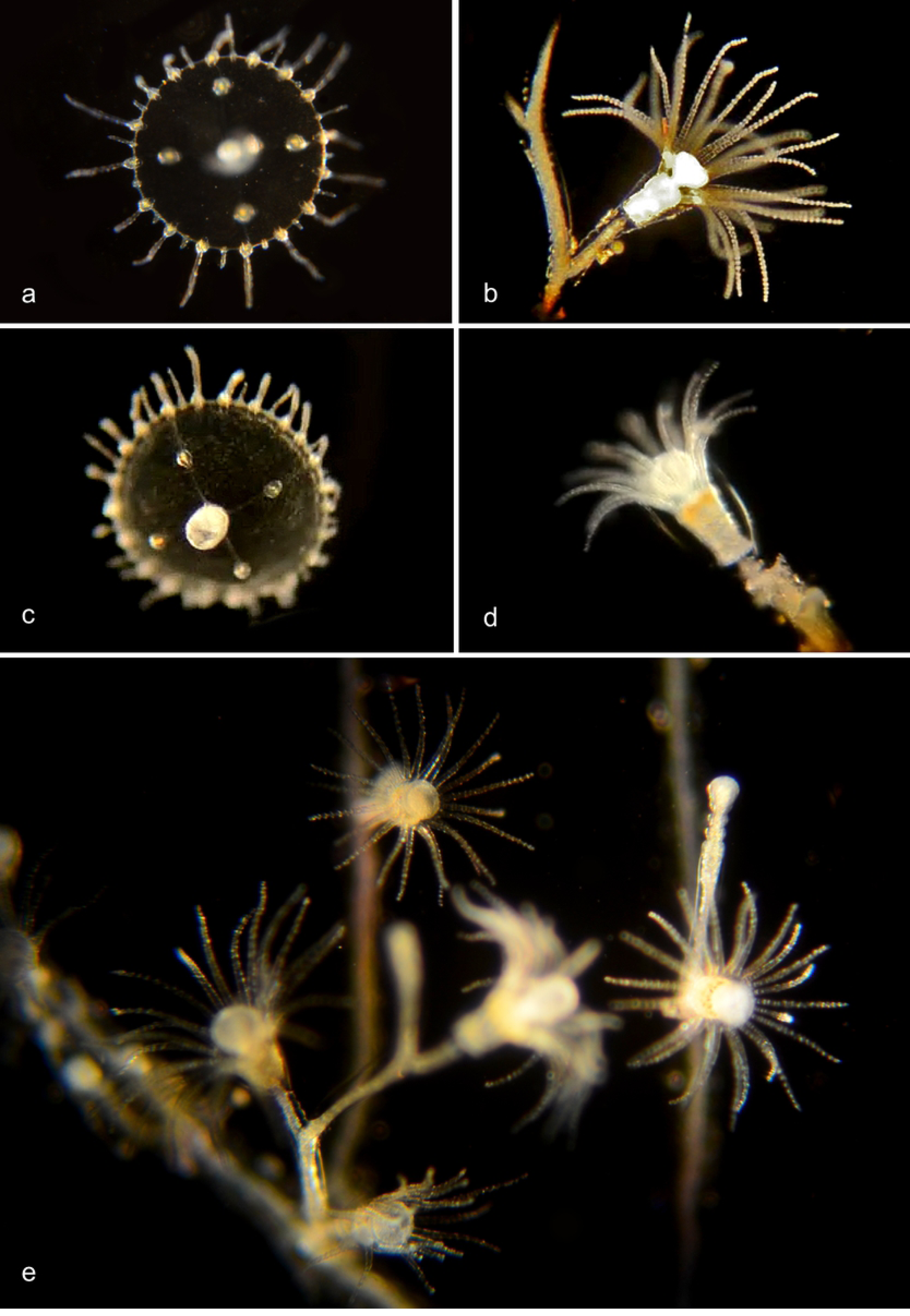
Clytia species. a. *Clytia elsaeoswaldae* (BIN ACR4294). Material collected: 2 medusae (from Balboa/Coral) as well as three medusae from Bahia de los Angeles, Mexico. BIOUG19284-D07. b. *Clytia hemisphaerica* (BIN AAD5403). Material collected: One hydroid BIOUG01227-E11 from Balboa/Coral. c,d. *Clytia gracilis* (BIN AAD5401). Material collected: One medusa from Newport Pier, one hydroid from Reef Point, Crystal Cove State Park. c, Medusa BIOUG01213-D07; d, Hydroid BIOUG19289-C05. e,f. *Clytia sp. 701 AC* (BIN AAR9451). Medusa phase: Four tentacles, four developing tentacles, and eight statocysts. Material collected: 3 medusae from off Dana Point, one from Lido/Genoa; One hydroid from Balboa/Coral). e, Medusa BIOUG19284-C12; f, Hydroid BIOUG01213-G11.

###### Genus Obelia

According to Cornelius (1975) the medusae of this genus are indistinguishable, and over seventy species had been described from the hydroid stage between 1830 and 1948. Cornelius referred all of these specimens to only three nominal species: *O. bidentata, O. dichotoma*, and *O. geniculata*. The identification characters given by Cornelius cannot all be used on our records, since they refer to growth habit and substrate, which has not always been recorded, or to the morphology of the hydrothecal rim which is usually not visible in our unstained specimens. Our DNA barcoding data identify thee species (Fig 11), of which one is clearly *O. dichotoma* and one is *O. geniculata*. The third group (*Obelia* sp. 1PJB) is presumably *O. bidentata*, although we have not been able to confirm that from the identification characters.

**Fig 11.**
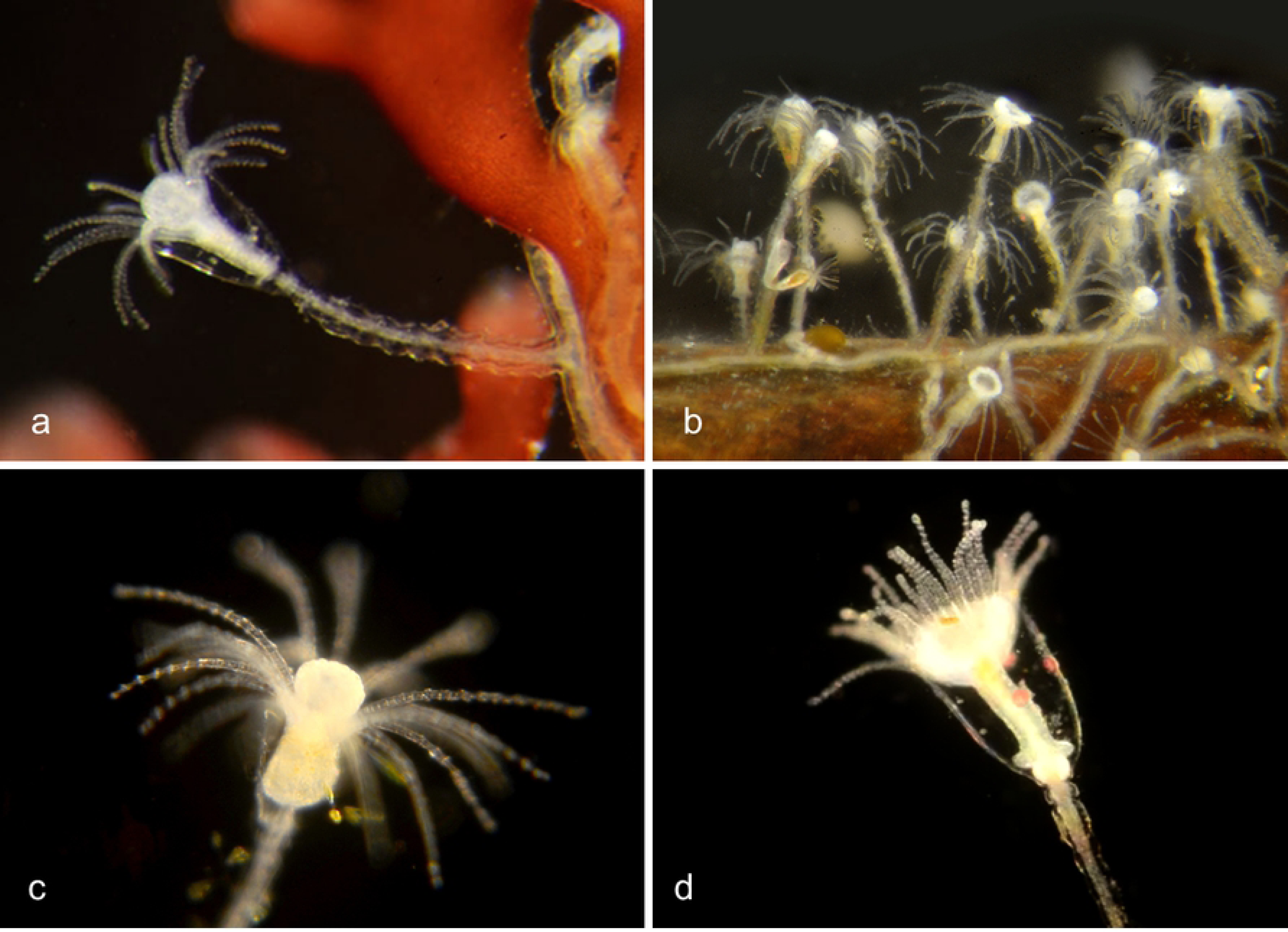
*Obelia* species. **a**,**b**. *Obelia dichotoma* (BIN ACO3910). Material collected: 5 medusae (3 from Newport Pier, one from Balboa/Coral, one from BBSC); 6 hydroids (5 from Newport Pier, one from Reef Point, CCSP). a, Medusa CCDB-25433-B12; b, Hydroid CCDB-24005-A10.

*Obelia sp*. 1PJB (BIN ACR4187). Material collected: 15 medusae (2 offshore from CCSP, 2 at NAC, 7 offshore from Dana Point, 2 from Harbor entrance, 1 from Balboa/Coral and one from PCH bridge); 10 hydroids (4 from Newport Pier, 5 from Balboa/Coral, one from Harbor). c, Medusa BIOUG012213-E12; d, Hydroid BIOUG01213-G10.

*Obelia geniculata* (BIN AAA708). Material collected: One hydroid, off Newport Pier. e, Hydroid BIOUG19284-G10

Genus *Orthopyxis* (Cunha et al, 2015)

Pedicel is annulated and has a smooth rim. We have not found medusa for any of these species.

**Figure 12.**
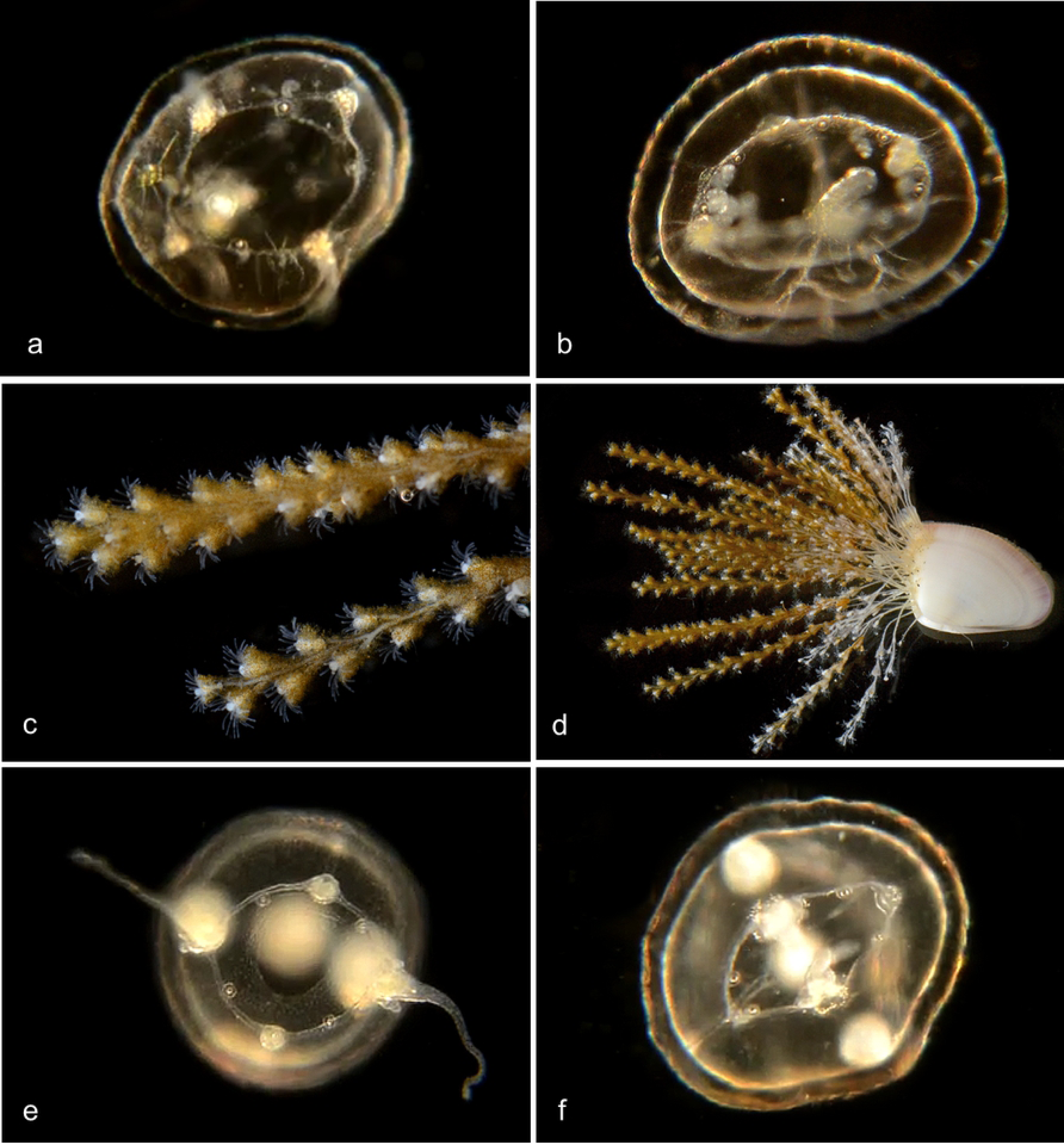
*Orthopyxis*

*Orthopyxis everta* (BIN ADE2083). Material collected: One hydroid from Reef Point, Crystal cove State Park. Hydroid BIOUG19289-E01.

*Orthopyxis sp*. 1PJB (BIN ADE2082). Material collected: One hydroid from Mussel Rocks, Crystal Cove State Park Hydroid BIOUG19289-D09.

*Orthopyxis sp*. 2PJB (BIN ACZ1022). Material collected: One hydroid from Newport Pier. Hydroid BIOUG19284-E01.

*Orthopyxis sp*. 3PJB (BIN ADH3873). Material collected: One hydroid from ocean off Newport Beach. Hydroid CCDB-25431-G09.

##### Family Lovenellidae

**Figure 13.**
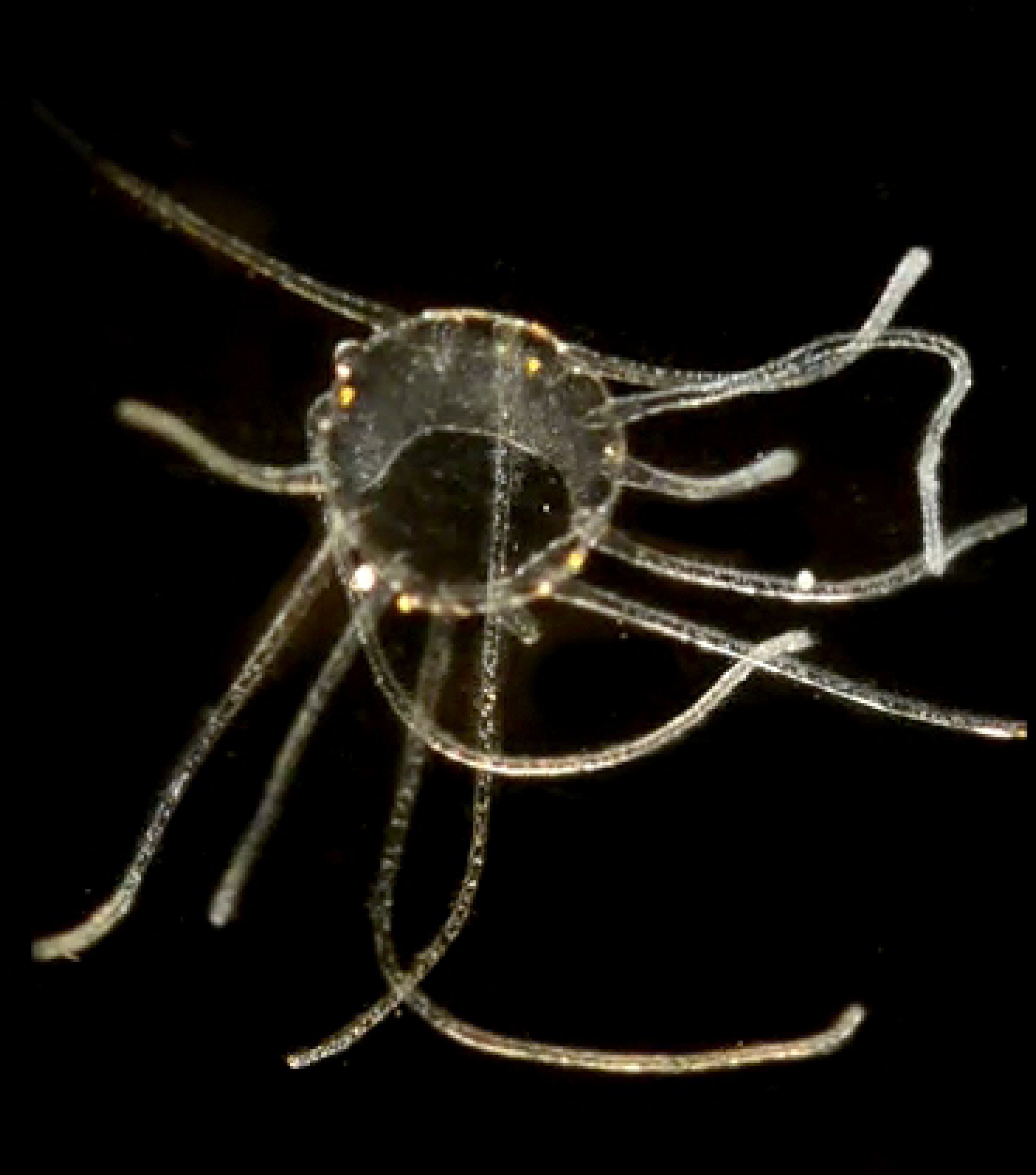
Genus *Eucheilota*. a. Close to *Eucheilota bakeri* (BIN ACW 1912 Torrey, 1909). 4 statocysts. Material collected: 5 medusae (2 at Newport Pier, 1 at Newport Aquatic Center, 2 at Balboa/Coral). Medusa BIOUG19284-D02 (Ventral). b. *Eucheilota sp*. (BIN ACR4297). 8 statocysts. Material collected: 3 medusae (1 at Crystal Cove, 2 at Newport Pier). Medusa BIOUG01227-E06. c-f. *Eucheilota bakeri* (BIN ACR4560) (Torrey, 1909). Material collected: Two hydroid colonies, attached to Bean Clam, *Donax gouldii*, washed up on the beach under Newport Pier. 4 statocysts. Two medusae, one from the Harbor entrance, one from Balboa/Coral. c, Hydroid BIOUG0213 C11; d, Hydroid BIOUG0213 C11; e, Medusa CCDB 24005 C03: f, Medusa CCDB-25431-H05.

#### Order Narcomedusae

Usually reproduce sexually as medusae, and do not form hydroids.

##### Family Solmarisidae (Schuchert, 2019a)

**Figure 14.**
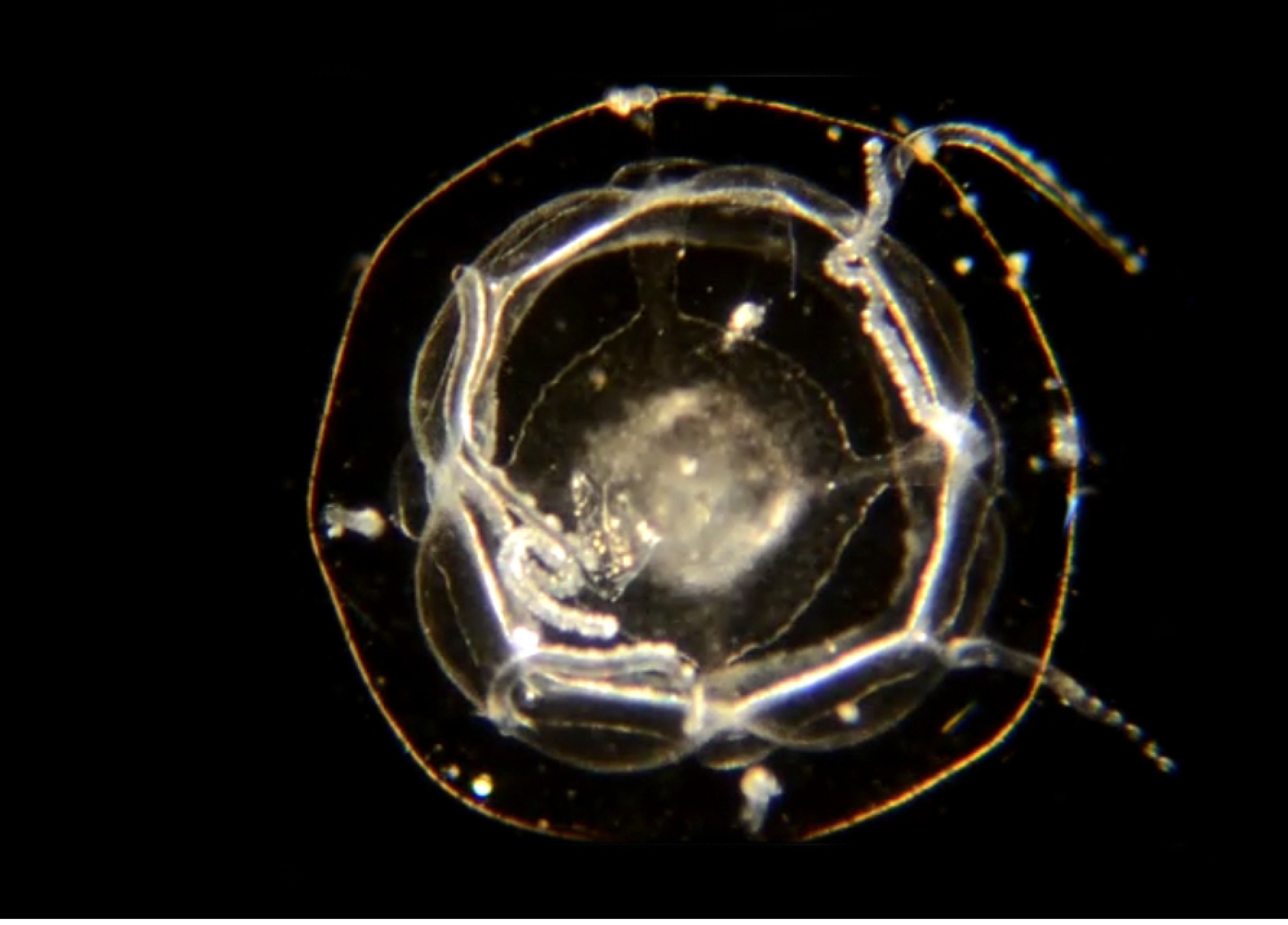
Medusa CCDB-25431-D07. Material collected: One medusa from Newport Harbor (not DNA barcoded).

#### Order Trachymedusae

Reproduce sexually as medusae, and do not form hydroids.

##### Family Geryoniidae

**Figure 15.**
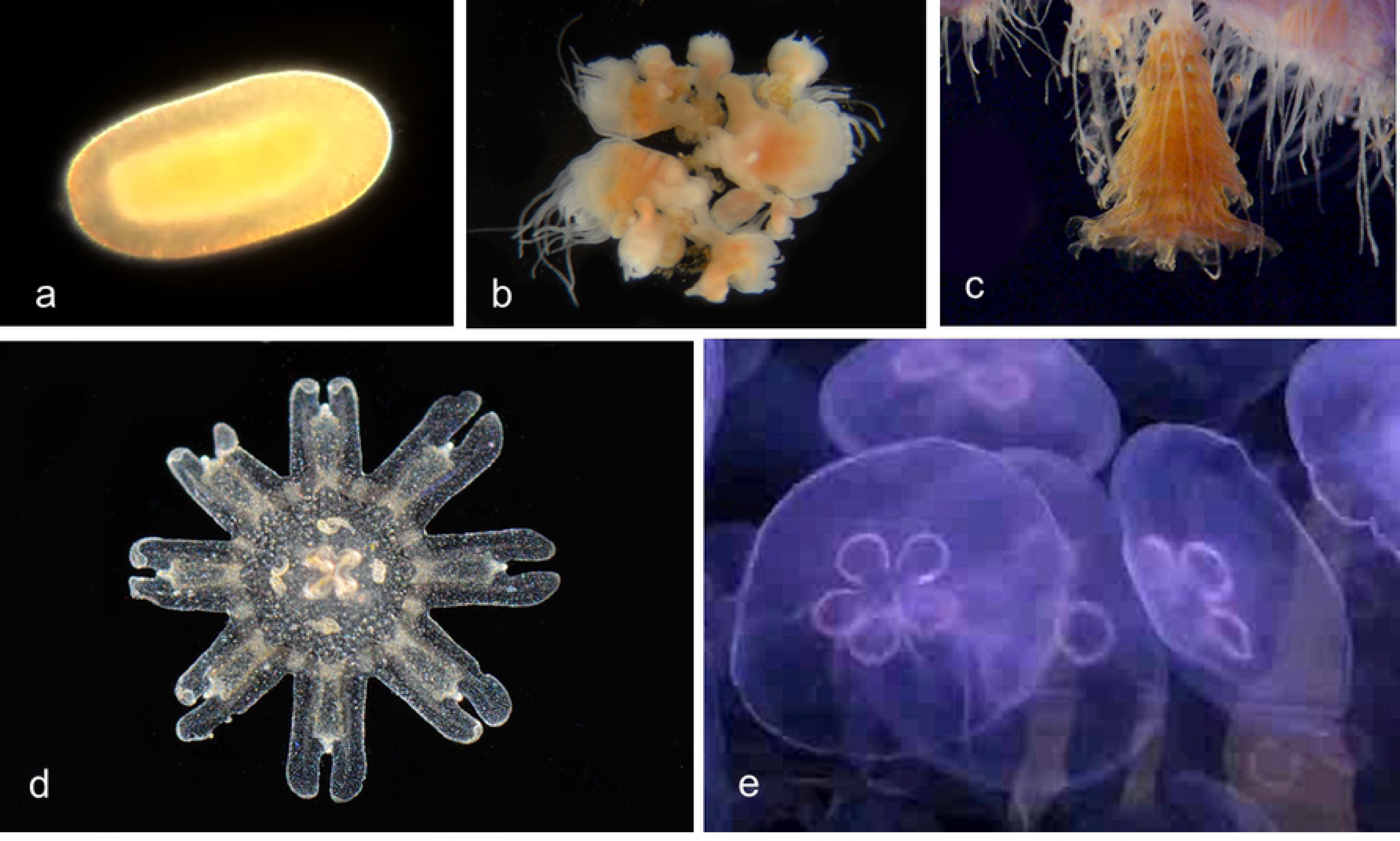
*Liriope tetraphylla*: BIN ACH9275. (Schuchert, 2019b). Material collected: 9 medusae (4 off Newport Pier, 2 off Dana Point, and one each from Balboa/Coral, Newport Harbor, and off CCSP). Medusa BIOUG01227-A02.

### Class Scyphozoa

The Scyphozoa are the true sea jellies, a name that refers to the sexually reproducing adult medusae. Fertilized eggs develop into planula larvae, which then develop into the colonies of asexually reproducing hydroids. The hydroid stage produces ephyra larvae by budding, and these larvae grow into the adult sea jellies. We have found only one species in our collections:

#### Order Semaeostomae; Family Ulmaridae

**Figure 16.**
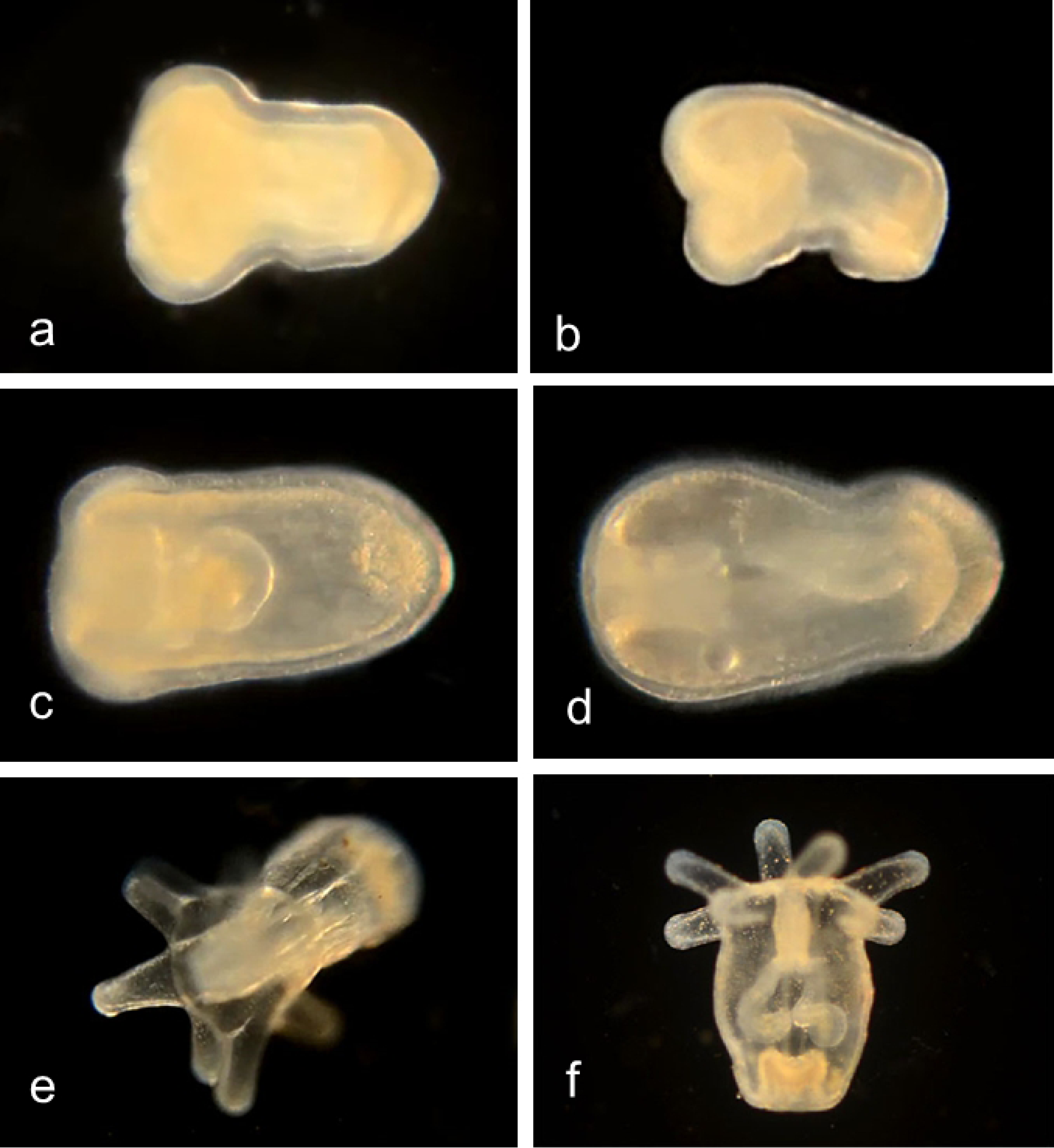
Moon Jelly, *Aurelia aurita* (BIN AAA4674) (Dawson, 2008). a, Planula larva (Material collected: one specimen from an adult collected off Lido Island; not DNA barcoded); b, Young hydroids (Material collected: 4 specimens from Cabrillo Marine Aquarium) BIOUG19284-A09. c, Strobilating hydroid (not DNA barcoded) from the Ocean Institute, Dana Point, CA. d, Ephyra larva BIOUG01213-C09 (Material Collected: 9 specimens from Lido/Genoa, 2 from NAC, 1 from Crew Dock); e, Adults (not DNA barcoded) at the Cabrillo Marine Aquarium, San Pedro, CA.

### Class Anthozoa

The Anthozoa includes corals, sea anemones, sea pens and seafans, in ten orders. Sea anemones have two types of larva that are found (rarely) in plankton samples: Antipathula and Cerinula. Our samples have included one of each larval type, from different species:

#### Order Actiniaria

**Figure 17.**
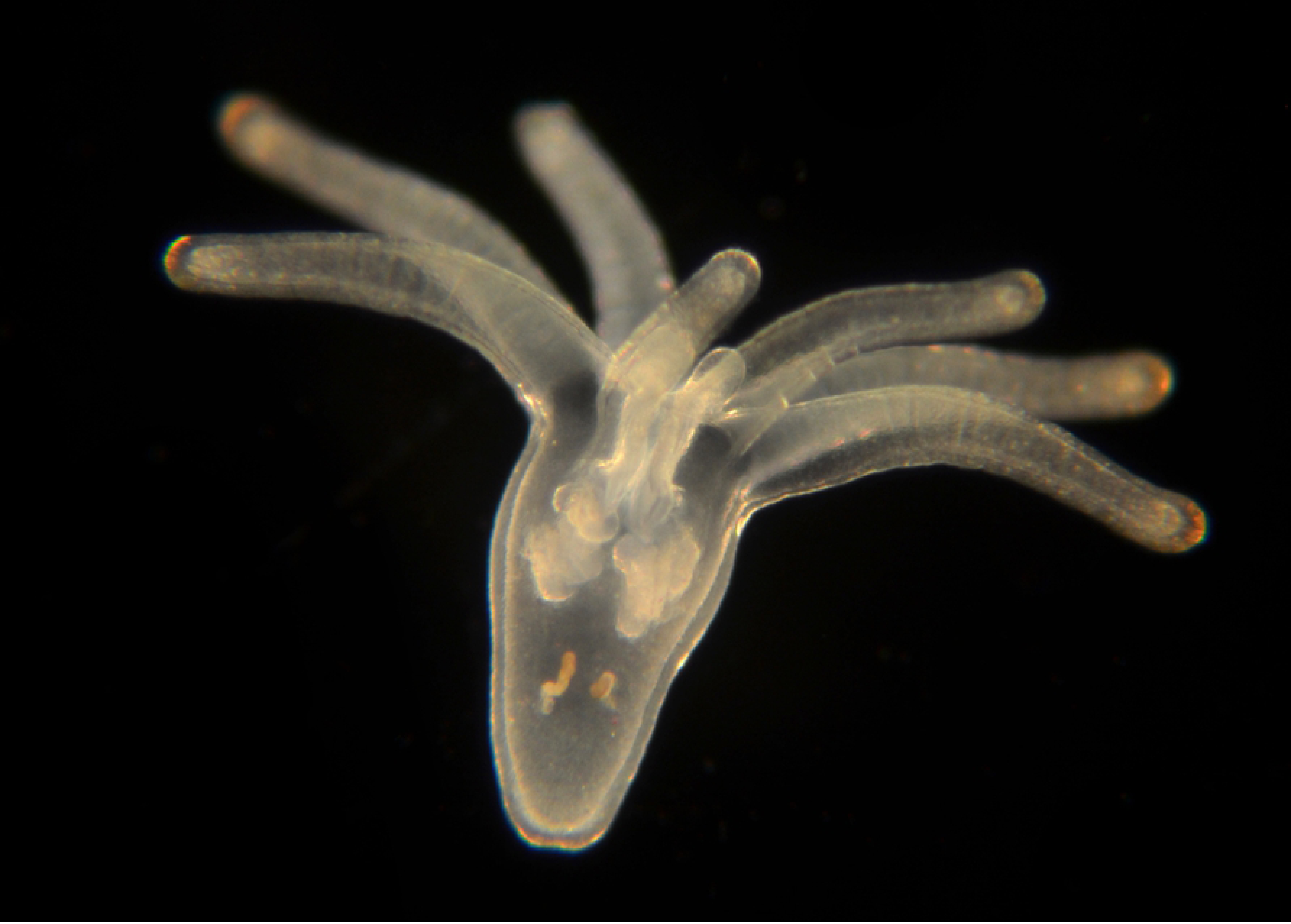
Onion anemone, *Paranthus rapiformis* (BIN ACC3637). Family Actinostolidae (Daly and Fautin, 2019). Material collected: Antipathula larvae: 2 from Newport Pier, 2 from Harbor entrance, two from off Newport Beach. a, CCDB-24005-F12; b, CCDB-24005-F11; c, CCDB-24005-H03; d, CCDB-24005-C01; e, BOIUG01213-F03; f, BIOUG19284-H08.

#### Order Penicillaria

**Figure 18.**
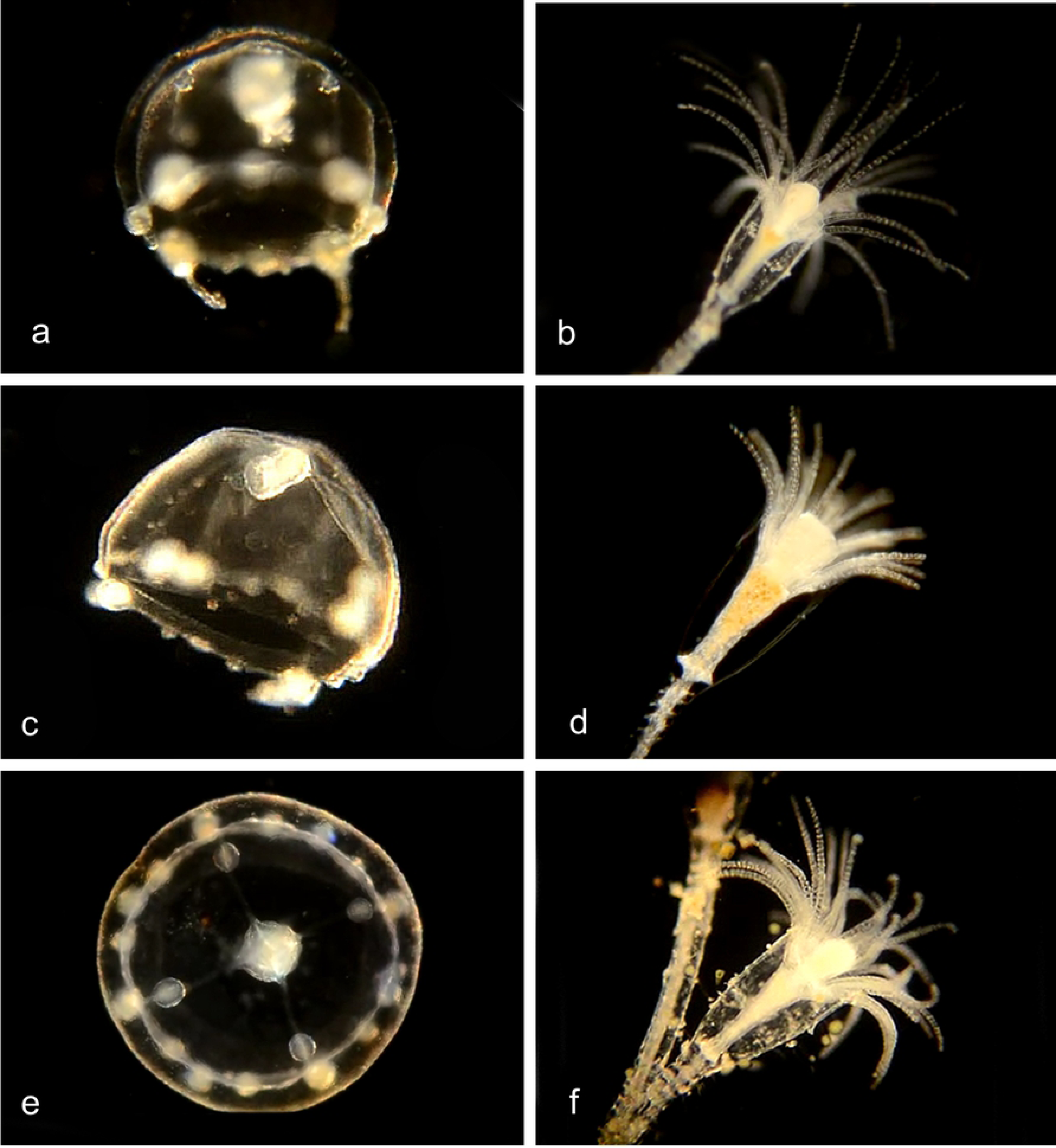
Tube-dwelling Anemone, *Isarachnanthus nocturnus* (BIN AAZ9217). Family Arachnactidae (Molodtsova, 2019). Cerinula larva. Material collected: One specimen BIOUG01213-B04 from off Newport Pier (and four from Baja California).

## Discussion

Our plankton samples, identified by DNA barcoding as well as morphology, have identified planktonic representatives of the classes Hydrozoa, Scyphozoa and Anthozoa, but no representatives of the class Cubozoa. One Cubozoan, *Carybdea confusa*, has been collected from Californian waters (Straehler-Pohl et al., 2017).

Among the order Anthomedusae (athecate hydroids), DNA barcoding allowed for the discrimination between the medusae of seven putative species of *Bougainvillia*. Of these, three species had medusae with four single tentacles, three (including the previously described *B. muscus*) with four paired tentacles, and two with four triplets of tentacles. The hydroid stages were documented for two of the *Bougainvillia* species.

We documented the medusa (with two simple tentacles) of three putative species of *Amphinema*, (including the previously described *A. dinema*) and documented the hydroid stages for two of them. DNA barcodes were obtained for medusae of one species of *Cladonema*, five putative species of *Corymorpha* (including the previously described *C. bigelowi)* with the matching hydroid phase for one; and *Coryne eximia, Turritopsis dohrnii* and *Turritopsis nutricula* with the matching hydroid phases for all of them. We identified the hydroid phase of *Coryne uchidai* by its DNA barcode. The hydroid and actinula larvae for the pink-hearted hydroid *Ectopleura crocea* were identified and matched by DNA barcoding.

Among the order Leptomedusae (thecate hydroids) medusae were identified for *Clytia elsaeoswaldae, Clytia gracilis* and *Clytia sp. 701 AC* and paired with the hydroid phases for the latter two species. Medusae were matched with the hydroid phases for two species of *Obelia* (including *O. dichotoma*) and *Eucheilota bakeri. O. geniculata* was collected as a single hydroid. Four species of *Orthopyxis*, (including *O. everta*), were collected only as hydroids.

One member of the family Solmarisidae, representing the order Narcomedusae, and one member (*Liriope tetraphylla*) of the order Trachymedusae were documented as medusae.

In the Scyphozoa, DNA barcoding confirmed the identification of the planktonic larval stage (ephyra) of the Moon Jelly, *Aurelia aurita*, the adult medusa of which is occasionally common in and around Newport Bay. We were also able to identify the planula stage (a free-swimming ciliated larva that eventually develops into the hydroid phase) for this species.

In the Anthozoa, antipathula larvae were identified from the Onion Anemone, *Paranthus rapiformis* and a cerinula larva was identified from the Tube-dwelling Anemone, *Isarachnanthus nocturnus*. We have yet to find the adults of these species locally.

The DNA differences in the COI barcode are, of course, probably not responsible for the morphological differences we have observed between specimens in separate taxa. However, the DNA differences between morphologically distinct medusae confirm that the specimens represent different taxa rather than different developmental stages within species. In some cases (e.g. *Bougainvillia, Amphinema, Liriope tetraphylla, Obelia*) DNA barcoding suggests that currently recognized species may include unrecognized distinct taxa, that are not obviously different at the morphological level.

The use of the DNA Barcode allows matching of hitherto unrecognized life-cycle stages within individual species, avoiding the ambiguities that have previously led to medusa and hydroid stages being assigned to separate species. Hydroid and medusa phases were originally matched by “circumstantial evidence” (Van Beneden, 1843; Cornelius, 1977) and later by culturing hydroid phases in the laboratory and observing the release of medusae (Russell, 1954, 1970). DNA barcoding is much less laborious than laboratory rearing, allowing for analysis on many more species. In the present study we have unambiguously matched medusa and hydroid phases for 13 species.

DNA barcoding has often revealed unexpected species diversity in many taxa (Waugh, 2007), and this study leads to the same conclusion in the realm of microscopic marine animals. It shows the utility of this approach and the value of the COI Barcode for identifying cnidarian species, in spite of published arguments against its use for this phylum (Hebert et al., 2003, Schuchert, 2010).

## Supplementary Materials

Bougainvillia tree

Amphinema tree

Cladonema tree

Corymorpha tree

Coryne tree

Turritopsis tree

Ectopleura tree

Clytia tree

Obelia tree

Orthopyxis tree

Campanulariidae tree

Eucheilota tree

Liriope tree

Aurelia tree

## Literature Cited

van Beneden, P. J. (1843). Mémoire sur les campanulaires de la côte d’Ostende considérés sous le rapport physiologique, embryogénique et zoologique. Ann. Sci. Nat. (Zool.) (2)20: 350–369.

Brinckmann-Voss (1970). Anthomedusae/Athecatae (Hydrozoa, Cnidaria) of the Mediterranean. Part I. Capitata. Fauna e Flora del Golfo di Napoli. 39. Stazione Zoologica. pp. 1–96.

Cornelius, P.F.S. (1975). The hydroid species of Obelia (Coelenterata, Hydrozoa: Campanulariidae), with notes on the medusa stage. Bull. Brit. Mus. (Nat. Hist.) Zoology 28: 251–293.

Cornelius, P.F.S. (1982). Hydroids and medusae of the family Companulariidae recorded from the eastern north Atlantic, with a world synopsis of genera. Bull. Brit. Mus. (Nat. Hist.) 42, 37–148.

Cunha, A.F., Genzano, G.N., and Marques, A.C. (2015). Reassessment of morphological diagnostic characters and species boundaries requires taxonomical changes for the genus Orthopyxis L. Agassiz, 1862 (Campanulariidae, Hydrozoa) and some related campanulariids. PLoS One; 10:e0117553. doi: 10.1371/journal.pone.0117553.

Daly, M.; Fautin, D. (2019). World List of Actiniaria. Paranthus rapiformis (Le Sueur, 1817). Accessed through: World Register of Marine Species at: http://www.marinespecies.org/aphia.php?p=taxdetails&id=158254on2019-05-28

Dawson, M. N. “Aurelia species”. Retrieved 2008-08-12. http://www2.eve.ucdavis.edu/mndawson/tS/Org/JotQ/JotQ_03Oct.html

Fofonoff, P.W., Ruiz, G.M., Steves, B., Simkanin, C., and Carlton, J.T. (2019). National Exotic Marine and Estuarine Species Information System. http://invasions.si.edu/nemesis/. Access Date: 26-May-2019

He, J., Zheng, L., Zhang, W., Lin, Y., & Cao, W. (2015). Morphology and molecular analyses of a new *Clytia* species (Cnidaria: Hydrozoa: Campanulariidae) from the East China Sea. Journal of the Marine Biological Association of the United Kingdom, 95(2), 289–300. doi:10.1017/S0025315414000836

Hebert, P.D.N., Ratnasingham, S. and deWaard, J.R. (2003). Barcoding animal life: cytochrome c oxidase subunit 1 divergences among closely related species. Proc. Bio. Sci. 270 (Suppl. 1), S96–S99.

Hubert, N. and Hanner, R. (2015). DNA Barcoding, species delineation and taxonomy: a historical perspective. DNA Barcodes 3: 44–58.

Hyman, L. H. (1947). Two new hydromedusae from the California Coast. Trans. Am. Microsc. Soc. 66: 262–268.

Lindner, A., Govindarajan, A.F., and Migotto, A.E. (2011). Cryptic species, life cycles, and the phylogeny of Clytia (Cnidaria: Hydrozoa: Campanulariidae) Zootaxa 2980: 23–36.

Marine Species Identification Portal: Liriope tetraphylla. Retrieved 27 May, 2019.

Molodtsova, T. (2019). World List of Ceriantharia. Isarachnanthus nocturnus (Hartog, 1977). Accessed through: World Register of Marine Species at: http://www.marinespecies.org/aphia.php?p=taxdetails&id=411138on2019-05-28

Russell, F. S. (1954). The Medusae of the British Isles. v. 1. Anthomedusae, Leptomedusae, Limnomedusae, Trachymedusae and Narcomedusae. Cambridge University Press, New York

Russell, F. S. (1970). Medusae of the British Isles. v. 2. Pelagic Scyphozoa with a supplement to the first volume on Hydromedusae. Cambridge University Press, New York.

Sassaman, C. and Rees, J.T. (1978). The life cycle of *Corymorpha* (= *Euphysora) bigelowi* (Maas, 1905) and its significance in the systematics of corymorphid hydromedusae. Bio. Bull. 154: 485–496. https://doi.org/10.2307/1541074

Schuchert, P. 2001. Survey of the family Corynidae (Cnidaria, Hydrozoa). Revue Suisse de Zoologie 108: 739–878. (Corynidae)

Schuchert, P. 2004. Revision of the European athecate hydroids and their medusae (Hydrozoa, Cnidaria): Families Oceanidae and Pachycordylidae. Revue Suisse de Zoologie 111: 315–369. (Turritopsis)

Schuchert, P. 2006. The European athecate hydroids and their medusae (Hydrozoa, Cnidaria): Capitata Part 1. Revue suisse de Zoologie 113: 325–410. (Cladonema)

Schuchert, P. (2010). The European athecate hydroids and their medusae (Hydrozoa, Cnidaria): Capitata Part 2. Revue Suisse de Zoologie, 117, 337–555. (Ectopleura crocea, Corymorpha)

Schuchert, P. (2007). The European athecate hydroids and their medusae (Hydrozoa, Cnidaria): Filifera Part 2.Revue Suisse de Zoologie, 114, 195–396. https://doi.org/10.5962/bhl.part.80395. (Bougainvillia)

Schuchert, P. (2019a). World Hydrozoa Database. Solmarisidae Haeckel, 1879. Accessed through: World Register of Marine Species at: http://www.marinespecies.org/aphia.php?p=taxdetails&id=16848on2019-05-27

Schuchert, P. (2019b). World Hydrozoa Database. Liriope tetraphylla (Chamisso & Eysenhardt, 1821). Accessed through: World Register of Marine Species at: http://www.marinespecies.org/aphia.php?p=taxdetails&id=117568on2019-05-28

Straehler-Pohl, I., Matsumoto, G.I., and Acevedo M.J. (2017) Recognition of the Californian cubozoan population as a new species *Carybdea confusa n. sp.* (Cnidaria, Cubozoa, Carybdeida). Plankton Benthos Res 12: 129–138.

Torrey, H.B. (1909) The Leptomedusae of the San Diego Region. University of California Publications in Zoology 6: 11–31.

Waugh, J. (2007). DNA barcoding in animal species: progress, potential and pitfalls. BioEssays 29: 188–197.

